# Band-selective IR PRESS for brain tumor spectroscopy allows robust detection of lactate

**DOI:** 10.1101/2024.09.30.615849

**Authors:** Shun Kishimoto, Olga Kim, Jeeva Munasinghe, Otowa Yasunori, Yamashita Kota, Kazutoshi Yamamoto, W. Marston Linehan, Jing Wu, Murali C Krishna, Jeffrey R Brender

**Affiliations:** Radiation Biology Branch, Center for Cancer Research, National Cancer Institute, National Institutes of Health, Bethesda, MD 20892, USA; Urologic Oncology Branch, Center for Cancer Research, National Cancer Institute, National Institutes of Health, Bethesda, MD 20892, USA; Neuro-Oncology Branch, Center for Cancer Research, National Cancer Institute, National Institutes of Health, Bethesda, MD 20892, USA; Laboratory of Functional and Molecular Imaging, National Institute of Neurological Disorders and Stroke, National Institutes of Health, Bethesda, MD 20892, USA; Molecular Imaging Branch, National Cancer Institute, National Institutes of Health, Bethesda, MD 20892, USA

## Abstract

Lactate plays a critical role in the tumor microenvironment, driving tumor progression, metastasis, and immune evasion. Despite its importance, in vivo quantification of lactate using magnetic resonance spectroscopy (MRS) has faced challenges, primarily due to the overlapping lipid signal at 1.3 ppm. Current clinical practice employs a long echo time to exploit differences in T2 relaxation between lactate and lipids; however, this approach significantly suppresses signals from other metabolites. Lipid has a notably different T1 relaxation time than lactate and other metabolites, which may be exploited by an inversion recovery sequence to better distinguish them. However, this method has not found wide use because of the loss of signal in other metabolites. Here we introduce a selective inversion pulse with a short echo time MRS method (SPIR-PRESS), which mitigates this issue. In phantom experiments, SPIR-PRESS successfully suppressed lipid signals that could be misinterpreted as lactate in short TE PRESS spectra, while maintaining sensitivity to the full metabolite profile. SPIR-PRESS demonstrated superior performance in quantifying lactate compared to long echo time PRESS, with ∼ 60 % increase in sensitivity for lactate detection compared to conventional PRESS with a 288 ms TE. In a mouse glioma model, SPIR-PRESS clearly detected lactate and other key tumor metabolites (choline, creatine, NAA) in the tumor, which were not detectable in conventional long TE PRESS. These findings highlight SPIR-PRESS as a promising technique for improved lactate quantification and comprehensive metabolite profiling in tumor environments.

---

Lactate is a key metabolite in cancer metabolism and the tumor microenvironment. In normal cells, lactate production occurs during glycolysis under anaerobic conditions. However, cancer cells frequently exhibit aerobic glycolysis (the “Warburg effect”), leading to increased lactate production even in the presence of oxygen.(1) Lactate can serve as an energy source for cancer cells,(2-4) promoting tumor growth and metastasis.(5) Furthermore, lactate accumulation in the tumor microenvironment contributes to an acidic and immunosuppressive environment,(6) facilitating tumor progression and treatment resistance.(7) Consequently, lactate has been investigated as a potential biomarker for cancer diagnosis, prognosis, and treatment monitoring.(8)

Emerging evidence suggests lactate is not only a passive byproduct of the Warburg effect, (1) but an active participant in immune resistance, metastasis, and tumor growth. Beyond its metabolic function, lactate acts as a multifaceted signaling molecule that drives cancer progression through multiple mechanisms. Lactate can be used as an alternative energy source for cancer cells, supporting their proliferation and survival.(2-4) Lactate also facilitates tumor cell migration and metastasis, contributing to cancer dissemination.(5) In the tumor microenvironment, it creates an acidic, (6) radio-(7) and chemoresistant,(9) immunosuppressive environment, enabling cancer cells to evade immune surveillance and contributing to resistance to therapeutic intervention across multiple treatment modalities. Consequently, lactate has been investigated as a potential biomarker for cancer diagnosis, prognosis, and treatment monitoring. (8)

In clinical settings, lactate is routinely quantified *in vivo* using magnetic resonance spectroscopy (MRS). Despite the strong mechanistic rationale for its use, attempts to use the lactate signal from MRS as a prognostic biomarker for glioblastoma and other neoplasms have had mixed results. While some studies have reported a positive correlation with tumor grade, two large multicenter studies found only infrequent occurrence of lactate in long TE single voxel studies with no obvious correlation with tumor grade. (10, 11) This discrepancy arises from the ambiguous nature of the signal at 1.3 ppm. The broad lipid signal at 1.3 ppm often dominates the short echo time spectrum of CNS tumors and healthy tissue of other organs. In the brain, the mobile triglyceride lipid signal arises from two sources: mobile lipids within the tumor (12, 13) and subcutaneous fat from the skull. While the subcutaneous fat signal can be removed by outer volume suppression (14) or k-space spatial filtering, mobile lipids from micrometer sized lipid droplets within the tumor cannot be easily removed by these techniques.

The most common method in clinical practice to distinguish the lactate peak from the overlapping broad lipid signal at 1.3 ppm exploits the faster transverse relaxation of lipids compared to lactate. (15) This is achieved by using a long echo time (TE) in the PRESS localization sequence. At longer TEs, the lipid signal is rapidly attenuated while the lactate signal persists due to its longer T2 relaxation time. However, the use of a long TE comes at the cost of reduced overall signal-to-noise ratio (SNR) and suppression of signals from other metabolites with shorter T2 relaxation times. Typical TE values used for lactate detection are 144 ms or 288 ms. At TE=144 ms, the lactate signal is inverted due to J-coupling evolution, but can suffer from signal cancellation via J-modulation effects. The problem can be particularly severe at 3T, where the chemical shift difference between the methyl and the methine proton causes severe signal cancellation due to anomalous J-modulation. (16) Therefore, a TE of 288 ms is more commonly used to unambiguously detect the lactate peak, though at the expense of very low SNR where the lactate signal may be the only discernible metabolite resonance remaining. An ideal solution would be a method capable of resolving the lactate and lipid signals at a short TE, maintaining sensitivity to the full metabolite profile while avoiding excessive signal losses.

To overcome the limitations associated with long TE sequences that rely on T2 differences, the substantial difference in T1 relaxation times between free lipid and lactate can be leveraged by using an inversion pulse preceding voxel spectroscopy to distinguish between the two metabolites, although no rigorous quantification or comparison with other methods has been performed to our knowledge. By using a spectrally selective inversion pulse to minimize T1 relaxation of other metabolites, we show that the short echo SPIR-PRESS and significantly outperforms the standard long echo (144/288 ms) sequence used in clinical practice for lactate quantification and allows the quantification of other metabolites.

## Results

The SPIR-PRESS pulse sequence is shown in Figure 1 and consists of a spectrally selective 180° inversion gaussian pulse inserted before a standard PRESS or ISIS pulse sequence with VAPOR or CHESS water suppression. The delay time after the inversion pulse (τ) is set to nullify the lipid signal. Relaxation after inversion occurs during the water suppression period, shortening the acquisition time considerably. A short echo time of 16.5 ms is used to maximize the lactate signal.

**Figure 1.**
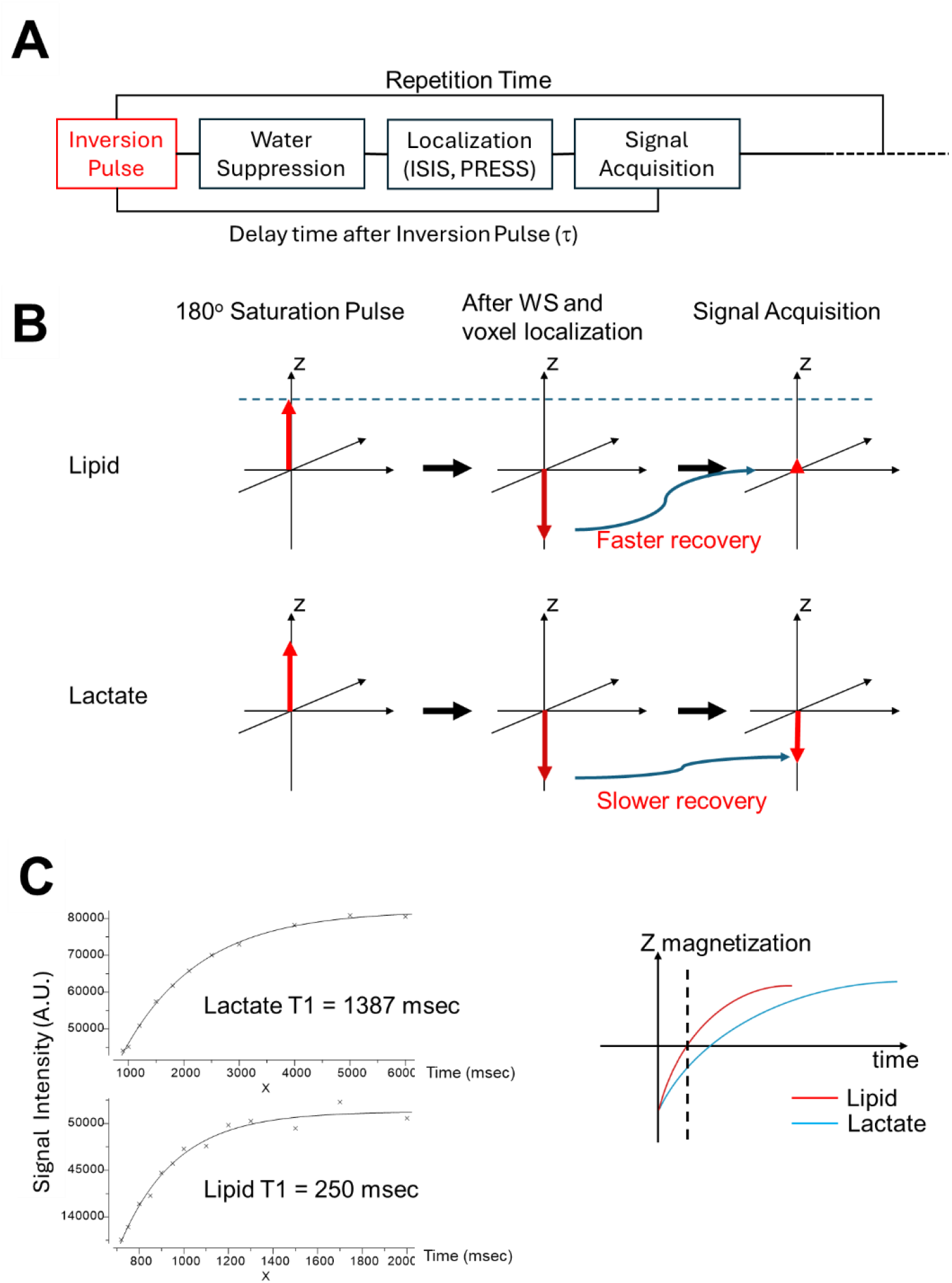
**A** Pulse sequence diagram of SPIR-PRESS. **B** Simplified evolution of the magnetization under SPIR-PRESS showing the nulling of the lipid single **C** T1 relaxation of a 111 mM lactate phantom (top) and 4% w/v lipid phantom (bottom)

The T1 nullification method requires a substantial difference in the T1 relaxation time between lipid and lactate. To measure the relaxation difference in a controlled environment, we created two one component phantoms: a 4% w/v lipid phantom (Intralipid, 30% Soybean Oil, 1.2% Egg Yolk Phospholipids, 1.7% Glycerin) and a 1% w/v lactate phantom pH neutralized through titration with sodium hydroxide. The T1 values of lipid and lactate, determined through a three-parameter exponential fitting are shown in Figure 2. The T1 of lactate (1387 msec) is more than five times longer than that of lipid (250 msec), potentially enabling the effective detection of negative z magnetization of lactate when the delay time after the inversion pulse (τ) is set to nullify the lipid signal (Fig. 1C and 1D).

**Figure 2.**
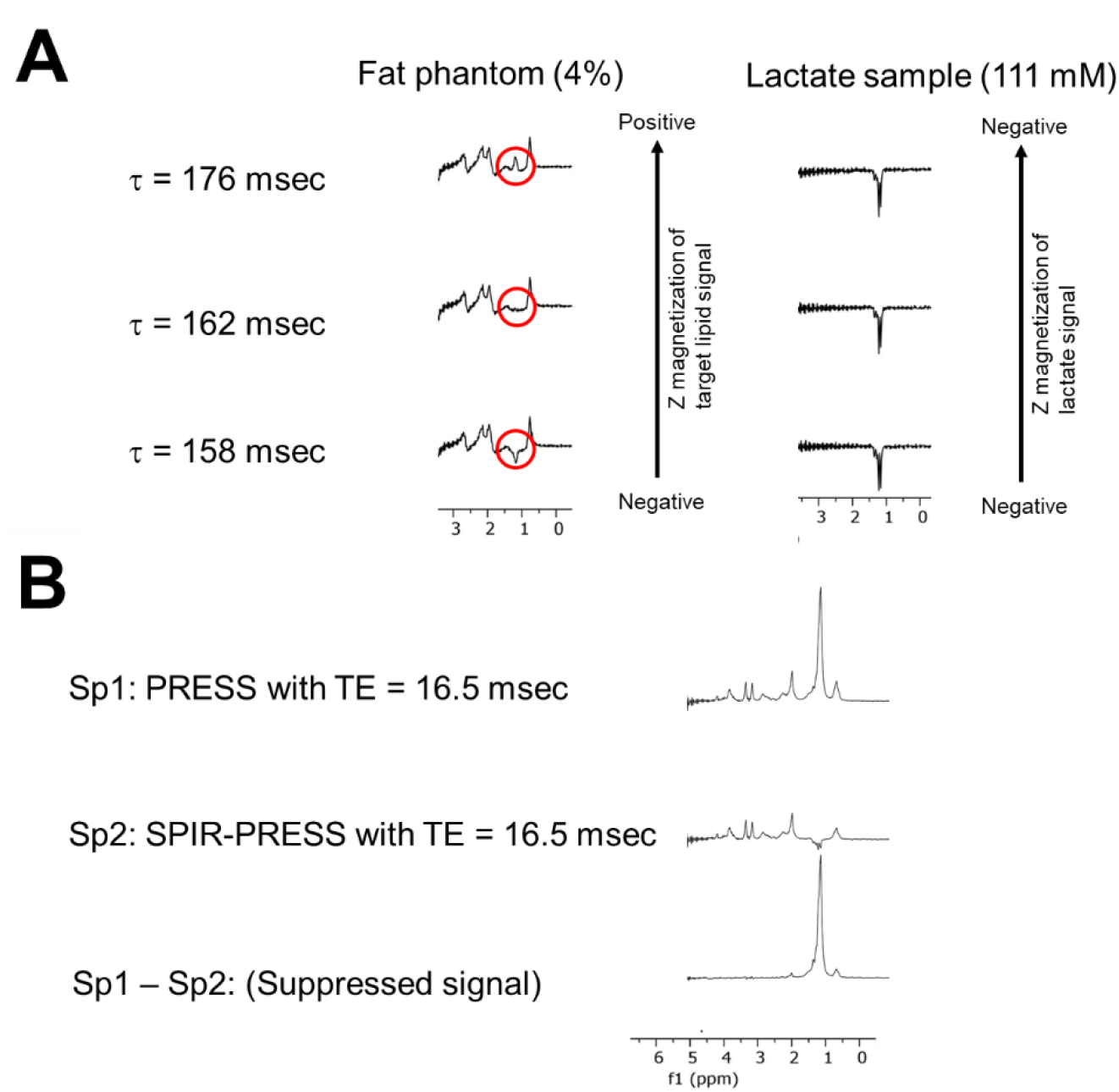
**A** Optimal null point of 4% Intralipid sample **B** 16.5 ms TE PRESS spectrum of a 4% Intralipid / 111 mM lactate sample (top) along with the corresponding SPIR-PRESS (middle) and difference spectra (bottom)

To characterize the lipid nulling effect, we analyzed SPIR-PRESS spectra from the 4% lipid phantom across a range of τ values (Figure 2A). The results show a clear progression from a negative lipid signal (158 msec) through a null point (162 msec) to a positive signal (176 msec), effectively mapping the inversion recovery process and identifying the optimal delay for lipid suppression. (Figure 2A). The potential advantages of SPIR-PRESS over conventional PRESS for lactate quantification are evident in the spectral comparison in Figure 2B. While the PRESS spectrum (bottom row) shows a dominant lipid peak that masks the lactate signal, SPIR-PRESS (middle row) successfully suppresses the lipid, exposing the lactate doublet at 1.32 ppm. The suppressed signal spectrum (Sp1 - Sp2) shows that the inversion pulse’s effect is limited to the 0.7-1.9 ppm range, underscoring the targeted nature of the band-selective approach.

We next assessed the quantitative performance of SPIR-PRESS in differentiating lactate signals from overlapping lipid signals. Phantoms containing lactate and 4% Intralipid were used to compare the relative sensitivity of SPIR-PRESS and conventional PRESS with a long echo time (TE=288 ms). Short echo SPIR-PRESS was found to have ∼ 60% increase in sensitivity compared to conventional PRESS with a 288 ms TE (Fig 3). To further evaluate the effectiveness of SPIR-PRESS in a more physiologically relevant setting, we prepared 1:1 lactate/lipid phantoms containing lactate, NAA, glutamate, choline, and creatine at concentrations mimicking those found in the brain under normal and pathological conditions (Lactate (2-20 mM), NAA (5.4-20 mM), choline (1.35-5.35 mM), glutamine (1-4.6 mM), creatine(6-10 mM), pH 7).(17-19). The delta Akaike Information Criterion(20) with correction for small sample sizes (ΔAICc)(21) was used to assess whether the regression slopes quantification differed significantly. This approach allows us to compare the potential sensitivity of the two techniques while accounting for our limited sample size.

**Figure 3.**
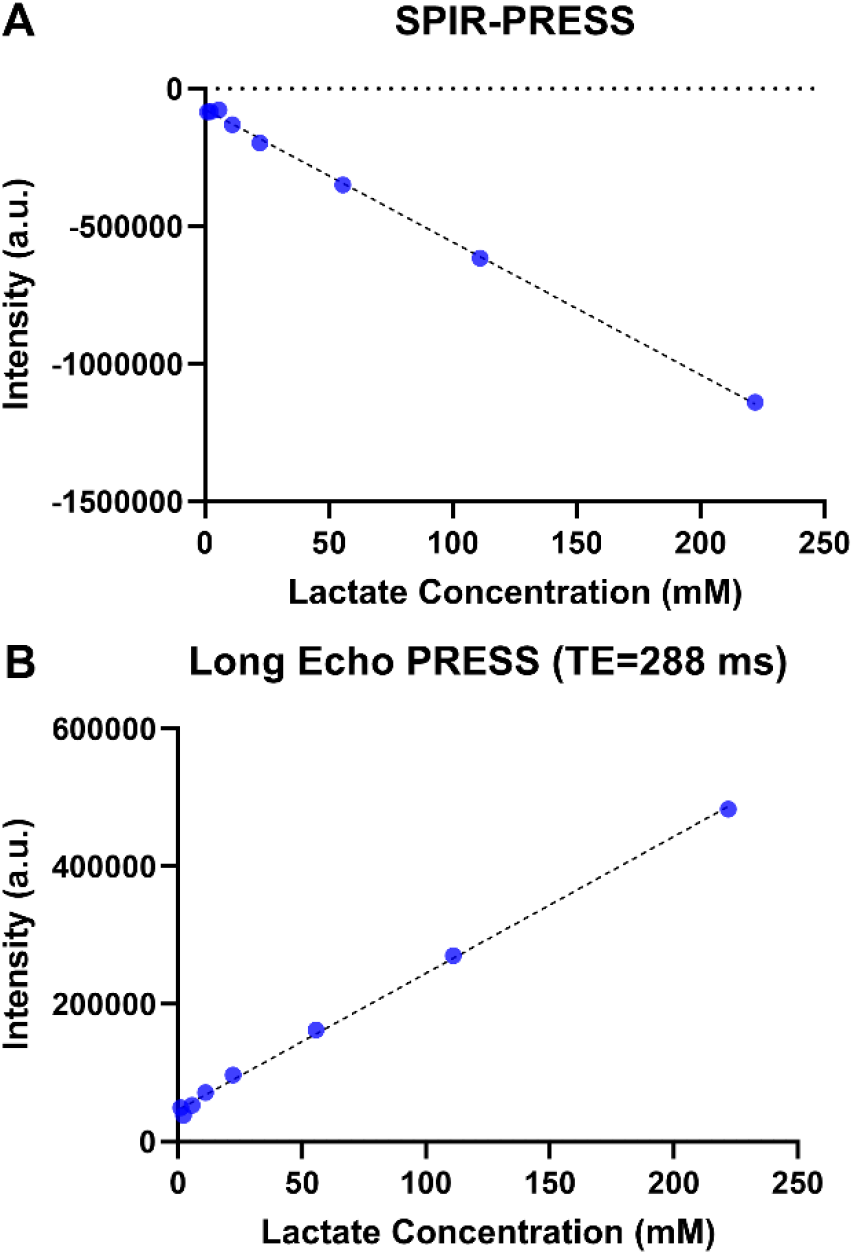
**A** Signal Intensity of the lactate peak in SPIR-PRESS spectra (TE=16.5 ms) of 4% Intralipid samples with varying concentrations of lactate. **B** Signal intensities of the lactate peak in long echo PRESS spectra of the same samples

As shown in Figure 4, the lactate peak was indistinguishable from the lipid signal when using PRESS with short echo time (TE=16 ms) or with intermediate echo time (TE=90 ms), which is more reflective of clinical practice. Although both long echo time PRESS (TE=288 ms) and SP-IRPRESS successfully quantified lactate with high accuracy (R>0.85), SPIR-PRESS demonstrated superior performance with a higher correlation coefficient (0.9382 vs. 0.8546). SPIR-PRESS also exhibited markedly higher sensitivity, as indicated by a significantly steeper regression slope when compared using delta ΔAICc (ΔAICc = 28.26, p<0.0001), likely due to its shorter echo time. For NAA, the metabolite peak closest to the inversion pulse, both methods showed equivalent sensitivity (ΔAICc = -3.13). The accuracy of NAA quantitation was significantly reduced when using PRESS with short and intermediate echo times, likely due to the presence of an unsuppressed lipid signal from the allylic carbon (-CH2-CH=CH-) resonating near 2 ppm. (22)

**Figure 4.**
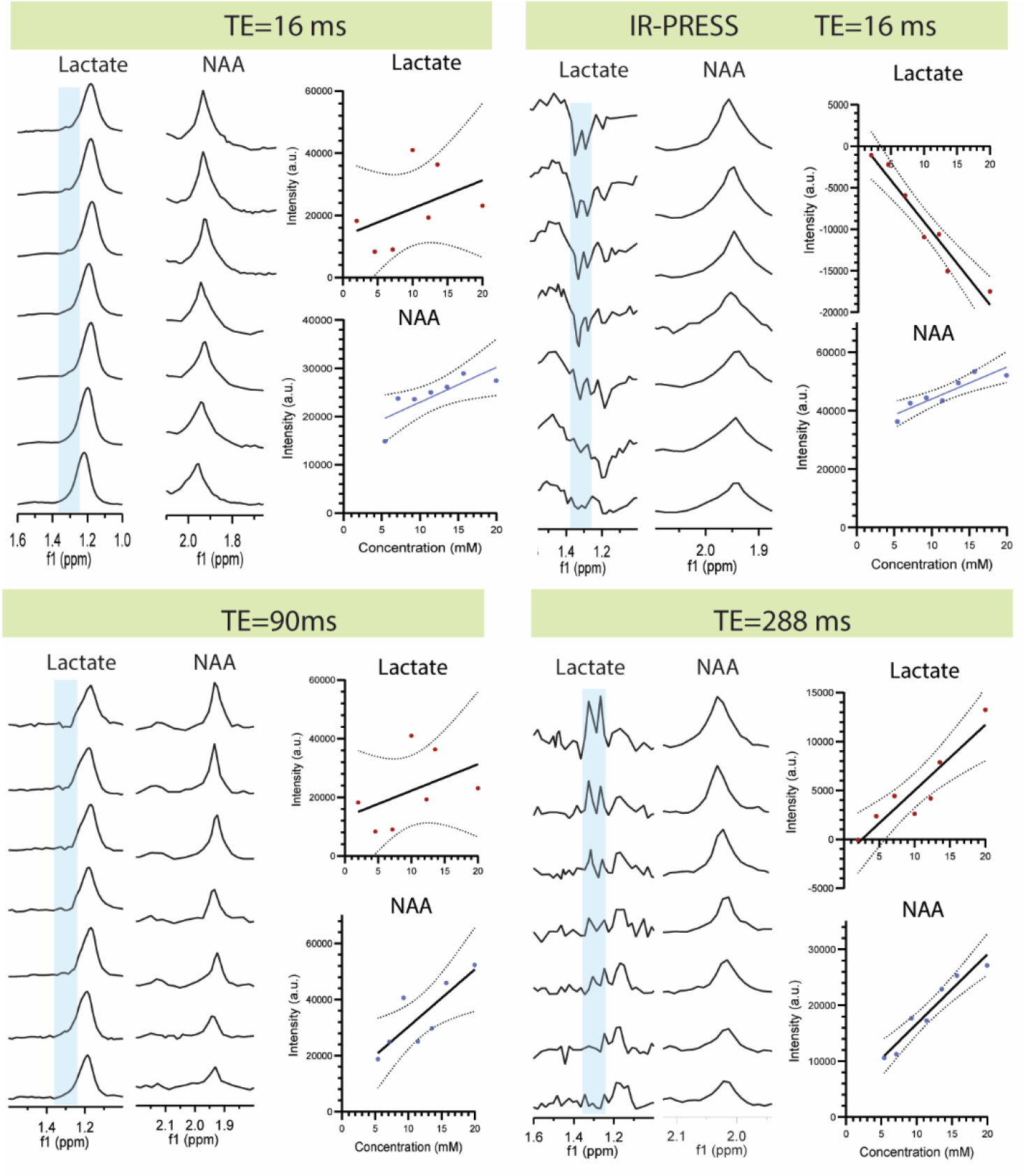
NAA and lactate quantification in a mixed phantom (Lactate (2-20 mM), NAA (5.4-20 mM), choline (1.35-5.35 mM), glutamine(1-4.6 mM), creatine(6-10 mM)) by SPIR-PRESS and PRESS with different echo times

We then investigated the impact of varying lipid content (0-16%) on lactate detection using PRESS and SPIR-PRESS sequences (Fig. 5). High concentrations of both lactate and lipid were used to detect any non-linearity in the measurements. Both methods clearly detected lactate, as evidenced by the doublet peaks in Fig. 5A, at a fixed lactate concentration of 1%. However, due to the shorter echo time (16.5 ms) used in SPIR-PRESS, some of the lipid signal was preserved, unlike in PRESS (288 ms), which completely eliminated the lipid signal. Consequently, SPIR-PRESS showed a constant lactate signal for lower lipid content (<6%), while PRESS maintained a constant signal across all lipid levels. This observation suggests that SPIR-PRESS may be susceptible to baseline shifts in the presence of strong lipid signals, potentially impacting the accuracy of lactate quantification. Despite this limitation, SPIR-PRESS demonstrated a significantly stronger lactate signal, more than twice that of PRESS, highlighting its potential for sensitive lactate detection in tissues with low to moderate fat content. However, caution should be exercised when using SPIR-PRESS in very fatty tissues, as the strong lipid signal may interfere with accurate lactate quantification.

**Figure 5.**
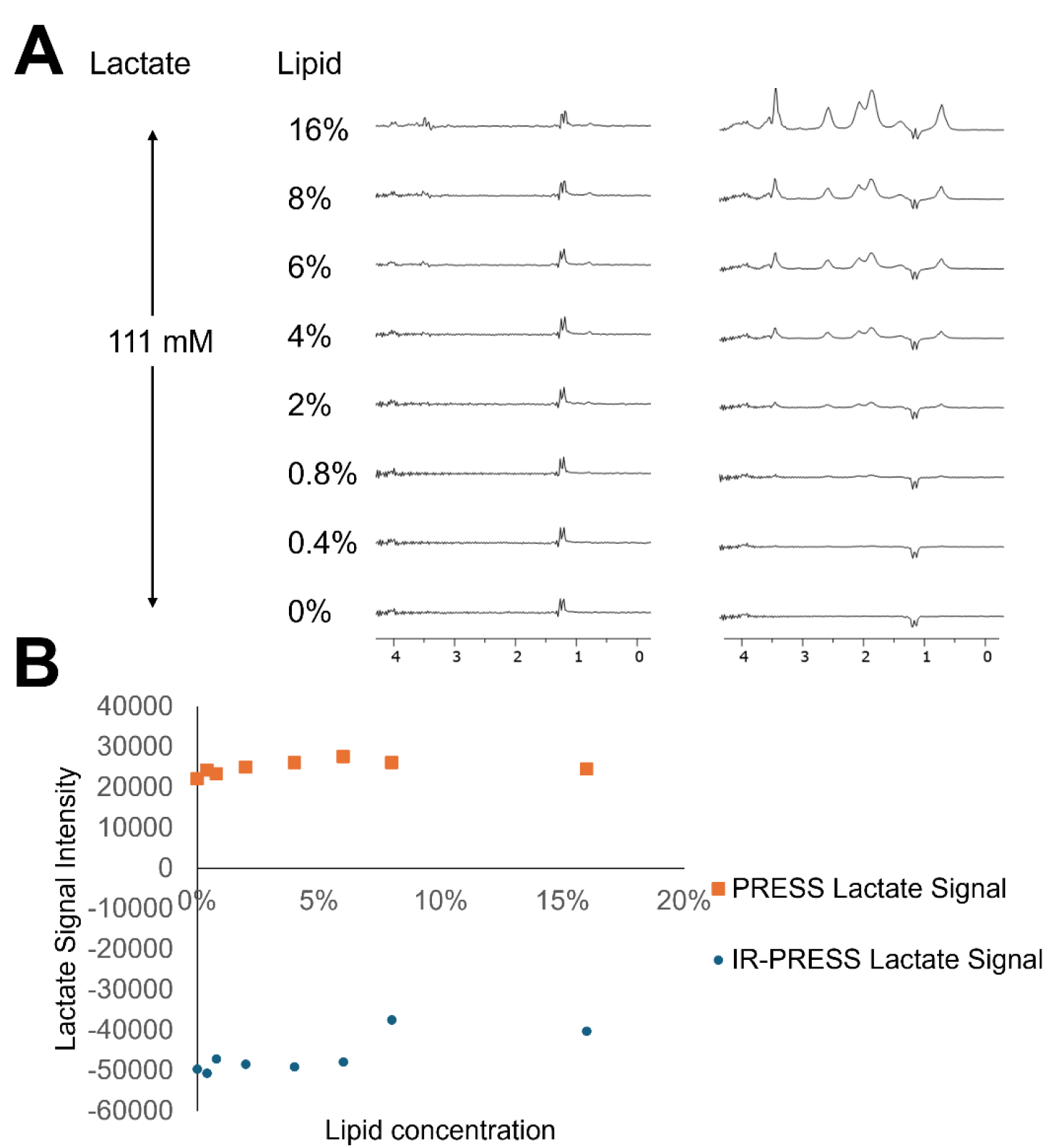
**A** Long Echo (TE=288ms) PRESS and short echo (TE=16.5) SPIR-PRESS specta of 111 mM lactate with varying lipid concentration. **B** Intensity plot of the lactate signal from both methods

To validate the SPIR-PRESS method in vivo, we compared its performance to that of conventional long TE PRESS in a mouse brain tumor model. 2 x 3 x 3 mm voxels were placed in the tumor and contralateral normal brain (Fig. 6A). The long TE (288 ms) PRESS tumor spectrum had poor sensitivity due to substantial T2 relaxation at long echo times, with only faint choline and creatine peaks visible (Fig. 6B). In contrast, the short TE (16.5 ms) SPIR-PRESS tumor spectrum clearly showed distinct choline, creatine, and NAA peaks. An inverted lactate peak at 1.3 ppm provided proof of the presence of lactate in the tumor (Fig. 5C, blue). Except for the negative lactate peak, the SPIR-PRESS spectrum is nearly identical to the short TE PRESS spectrum without the inversion pulse (blue), confirming that the measurement of other metabolites was not affected by the inversion pulse.

**Figure 6.**
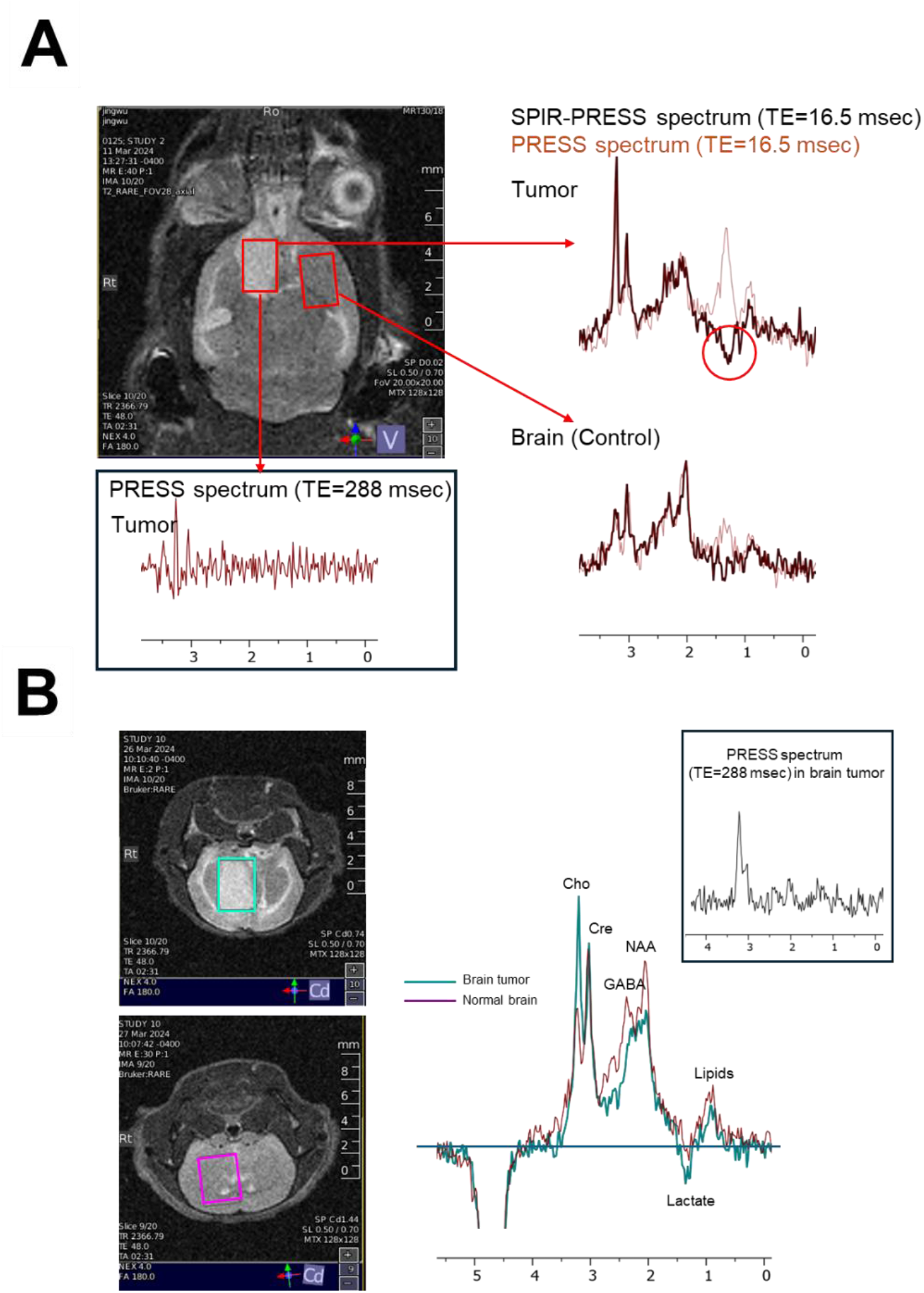
**A** Short echo PRESS (pink) and SPIR-PRESS (black) from a mouse glioma and the corresponding normal tissue B Comparison of long PRESS (inset) and short echo SPIR-PRESS within a mouse glioma

In the contralateral negative control voxel, a peak at 1.3 ppm is present in the short TE PRESS spectrum that could be confused for lactate. However, this peak is absent in the SPIR-PRESS spectrum, indicating it arises from lipids rather than lactate. Given that lactate in normal brain grey matter is only (∼2 μmol/g), this peak is almost certainly lipid, indicating successful nulling of the lipid signal at 1.3 ppm by the inversion pulse. These findings underscore the capability of SPIR-PRESS to accurately identify key tumor metabolites, including lactate, without interference from lipid signals that could be misinterpreted in the short TE PRESS spectrum. This represents a significant improvement over the conventional long TE PRESS, which failed to detect lactate and whose spectrum is predominantly characterized by noise.

## Discussion

We showed that the SPIR-PRESS method can effectively detect lactate in the presence of overlapping lipid signals at short echo times, providing a significant improvement over conventional long TE PRESS. Our approach is similar to that described by Balchandani et al. for fat suppression in 1H MRSI at 7T.(23) However, their method utilized an adiabatic inversion pulse with a longer duration (30 ms), sacrificing some SNR for improved B1 insensitivity. In contrast, we employ a shorter duration (11 ms), spectrally selective inversion pulse to minimize T1 relaxation of other metabolites while still achieving lipid nulling. This allows us to use a much shorter echo time (16.5 ms) compared to the standard long TE (144/288 ms) PRESS sequence used in clinical practice for lactate detection. By employing this band-selective inversion pulse, SPIR-PRESS exploits the substantial difference in T1 relaxation times between lactate and lipids to selectively null the lipid signal while preserving the lactate signal and other metabolite peaks. This approach maintains sensitivity to the full metabolite profile, which is crucial for diagnostic and monitoring purposes in brain tumors.

The phantom experiments demonstrate that SPIR-PRESS can accurately distinguish lactate from lipid signals, with a potentially lower limit of detection compared to conventional long TE PRESS. Furthermore, our *in vivo* results in a mouse brain tumor model confirm the ability of SPIR-PRESS to detect lactate in tumors while suppressing lipid signals that could be misinterpreted as lactate in short TE PRESS spectra. The clear detection of other key tumor metabolites, such as choline, creatine, and NAA, highlights the potential of SPIR-PRESS as a valuable tool for non-invasive assessment of brain tumor metabolism.

Spectral editing techniques have been utilized to resolve the lactate and lipid signals at a short TE,(24, 25) aiming to maintain sensitivity to the full metabolite profile while avoiding excessive signal losses. However, these methods rely on exact cancellation between scans, making them susceptible to motion artifacts that nullify the exact cancellation.(26) Multiple quantum filtering (MQF) has been proposed to address these concerns. However, MQF methods yield spectra that display only coupled spin systems, thereby eliminating other biologic markers for disease, such as N-acetylaspartic acid (NAA), creatine (Cr), and choline (Cho), which are valuable for diagnostic and monitoring purposes.

The limitations of SPIR-PRESS as implemented here are primarily those inherent to all short echo time spectroscopy techniques. Glutamate and glutamine in particular are not resolved from each other or resolved fully from the NAA peak. The selective pulse only suppresses part of the macromolecule baseline, which may cause problems in quantitative fitting. The narrow selective pulse used here does not invert the α-olefenic lipid peaks at 2.2 and 2.5 ppm, although the problem is exaggerated in the phantom by the very high content of PUFA in soybean oil (58%) compared to human tissue (2%).(27) Finally, the lactate signal is also attenuated slightly by T1 relaxation after the inversion pulse. Despite these limitations, SPIR-PRESS represents a promising approach for detecting lactate in brain tumors with improved sensitivity and specificity compared to conventional long TE PRESS, enabling more accurate assessment of the role of lactate in tumor metabolism and mitochondrial disease. Further validation in human studies is warranted to establish its clinical utility.

## Methods

### Animal study

The glioblastoma model mouse was prepared by using an athymic mouse inoculated with patient derived glioblastoma cell line GBM1 to obtain MRS of the tumor.(28) Eight-week-old female NSG mice were anesthetized with a combination of xylazine (20mg/ml) and ketamine (100mg/ml) diluted in 0.9% injection NaCl at a 1:1:4 volume ratios for a total dose of 0.1ml per 20g body weight. After the animals were fully anesthetized, they were placed and immobilized in a small animal stereotactic frame fitted with a mouse-specific headpiece using lidocaine gel pre-treated ear bars. All surgery procedures were done under sterile condition. GBM1 cells (0.5×10^6^ cells/2mL) resuspended in serum-free Neurobasal A medium were slowly injected intracranially (2 mm anterior and 2 mm lateral to bregma; 2.5 mm deep from the dura). The burr hole was closed with bone wax, and the scalp closed with Vetbond, or one to two staples or sutures.

Animals were observed frequently while they recover from the anesthesia in a warm, draft-free area. All animal experiments were approved by the National Cancer Institute Animal Care and Use Committee (NCI ACUC) and conducted in accordance with NCI ACUC guidelines under the authority of the animal protocol (NOB-023). During anesthesia, a pressure transducer (SA Instruments Inc.) was used to monitor the respiratory rate, which was maintained at 60 ± 10 breaths per minute. A nonmagnetic rectal temperature probe (FISO) was used to monitor the core body temperature, which was maintained at 36 ± 1°C using a circulating water-warming pad.

T2-weighted MRI images were acquired using a 3 T Biospec MRI scanner with a Rapid Acquisition with Relaxation Enhancement (RARE) pulse sequence. The following parameters were used: repetition time (TR) = 2366 ms, echo train length = 8, echo time (TE) = 48 ms, and number of averages = 4. The imaging volume consisted of 20 slices, each with a size of 2 cm × 2 cm and a thickness of 3 mm.

## Magnetic resonance spectroscopy

All spectroscopy data were acquired by custom made coil with 3 Tesla Bruker Biospec system. For phantom experiments, the following parameters were used for SPIR-PRESS sequence. TE = 16.5 msec, TR = 7000 msec, number of averages = 32, spectral width = 2564.1 Hz (20 ppm). Voxels of 8 x 8 x 8 mm were used for the experiments with phantoms of varying lipid compositions (Figure 3) and 5 x 5 x 5 mm voxels were used for other experiments. For all phantom studies, the pH of lactate was adjusted to 7.0 by titration with sodium hydroxide. The CHESS pulse sequence with an excitation width of 140 Hz was used for water suppression. For SPIR-PRESS, an 18 msec Gaussian inversion pulse with a bandwidth of 150 Hz was used. For conventional PRESS sequence experiments, the same setting was used except for an echo delay of TE=144 or 288 msec and no inversion pulse. TR = 7000 msec, number of averages = 8, spectral width = 2564.1 Hz (20 ppm), and voxel size of 5 x 5 x 5 mm were used for ISIS sequence.

For animal experiments, TR = 7000 msec, number of averages = 300, spectral width = 2564.1 Hz (20 ppm). Voxel size of 3 x 4 x 4 mm, CHESS water suppression sequence with a 350 Hz bandwifth, TE = 16.5 msec for SPIR-PRESS / TE = 288 msec for PRESS, and frequency selective inversion pulse (Gaussian) of 250 Hz (10.96 msec) were used for spectroscopy. The FID data were processed by Matlab software and MestreNova for quantification of the metabolite peaks. Linear regression was performed in Graphpad Prism 10.3.0. As the metabolic profile of the contralateral normal brain region could be influenced by the large tumor, another untreated athymic mouse was also prepared to obtain MRS of normal brain tissue control.

